# Medial prefrontal cortical NMDA receptors regulate depression-like behavior and dictate limbic thalamus innervation

**DOI:** 10.1101/106419

**Authors:** Oliver H. Miller, Andreas Bruns, Imen Ben Ammar, Thomas Mueggler, Benjamin J. Hall

## Abstract

Depression is a pervasive and debilitating neuropsychiatric disorder. A single, low dose of the NMDA receptor (NMDAR) antagonist ketamine elicits a long-lasting antidepressant response in patients with treatment-resistant major depressive disorder. Developing mechanistic understanding of how NMDAR antagonism alters synapse and circuit function is pivotal to developing translatable, circuit-based therapies for depression. Here using viral vectors, anatomical tracing, fMRI, and optogenetic-assisted circuit analysis, we assessed the role of the NMDAR subunit GluN2B in regulating cellular, synaptic, and circuit-level function and depression-related behavior. We demonstrate that post-developmental deletion of GluN2B from pyramidal neurons in medial prefrontal cortex enhances action potential output in a synaptic activity-dependent manner. GluN2B deletion dictates functional connectivity between mPFC and limbic thalamus but not ventral hippocampus and elicits antidepressant-like behavior. Our findings demonstrate that postsynaptic GluN2B exerts input-specific control of pyramidal neuron innervation, and identify a novel circuit for regulating depression-like behaviors in mice.

## Introduction

One in twenty people experience a depressive episode in their lifetime^1^. The limited efficacy of primary treatments and limited accessibility of efficacious treatments leave a large proportion of depressed patients untreated^2^. N-methyl D-aspartate receptors (NMDARs) represent an important and mechanistically distinct class of targets for treatment of major depressive disorder (MDD)^3–6^. A low dose of the NMDAR antagonist ketamine elicits antidepressant effects in treatment-resistant patients within 110 minutes, with this single dose remaining efficacious over placebo for at least 7 days^7^. While clinical work has broadened the scope of disorders targetable by NMDAR antagonism, pre-clinical studies are providing a mechanistic understanding of how it is that a brief period of NMDAR modulation has such powerful and sustained effects on mood. Recent work suggests that antagonism of GluN2B-containing NMDARs on forebrain pyramidal neurons is a sufficient and necessary component of ketamine’s antidepressant-like actions^8–11^. However, it is not clear which forebrain regions and brain circuits play a causal role in inducing the antidepressant-like effects of NMDAR antagonism.

The medial prefrontal cortex (mPFC) is involved in the motivational aspects of depression-related behavior in rodents and its activity is bi-directionally affected by the induction of depression or treatment with antidepressant drugs^12–14^. It has been demonstrated that a single treatment with low-dose ketamine or the GluN2B-subunit selective antagonist Ro 25-6981 results in an increase in excitatory input onto mPFC neurons in rodents coincident with the antidepressant-like response^9,11,14^. The distinct inputs giving rise to these strengthened synaptic connections and their role in altering mPFC circuit function, however, remains unknown.

In the current study, we demonstrate that post-developmental deletion of GluN2B from pyramidal neurons in mPFC is sufficient to elicit antidepressant-like behavior in mice. GluN2B deletion enhances excitatory synaptic drive onto layer 3 pyramidal neurons in mPFC, resulting in a synaptic activity-dependent facilitation of action potential output. Using optogenetic-assisted electrophysiology, we found that synaptic remodeling following ++GluN2B deletion occurs at local mPFC microcircuits engaged by inputs from the medial dorsal thalamus (MDT), but not the ventral hippocampus (vHipp). Finally, to observe alterations in activity of mPFC-connected nuclei *in vivo,* we employed arterial spin labeling (ASL)-based perfusion fMRI after mPFC GluN2B deletion and observed altered perfusion in mPFC and MDT. Together these data strongly implicate mPFC GluN2B-containing NMDARs in regulating depression-related behavior, and suggest that input-specific control over mPFC microcircuits may underlie such behavioral changes.

## Results

### Deletion of GluN2B from mPFC pyramidal neurons elicits antidepressant-like behavior

Selective, forebrain deletion of GluN2B from cortical pyramidal neurons is sufficient to recapitulate the antidepressant actions of ketamine. Application of ketamine results in no additional antidepressant effect in these mutant animals, suggesting that ketamine exerts its effects by antagonizing GluN2B-containing NMDARs in forebrain pyramidal neurons^9^. However, it is not clear which forebrain regions and circuits play a causal role inducing the antidepressant-like effects of ketamine.

The mPFC is a primary locus of top-down executive control over the processing of aversive and appetitive stimuli, and has a well-established role in the regulation of mood^15–18^. For these reasons we sought to determine if removal of GluN2B from pyramidal neurons in mPFC alone would be sufficient to elicit antidepressant-like effects. To specifically remove GluN2B from mPFC excitatory neurons, we targeted an adeno-associated virus (AAV) expressing Cre-GFP or a GFP control virus under the CaMKII promotor (AAV2-CaMKII-GFP-T2A-Cre or AAV2-CaMKII-GFP) bilaterally to the superficial layers of the prelimbic mPFC of GluN2B^floxed^ adult mice (2BΔ^mPFC^ mice, Figure 1A-C). Two weeks after injection, minimal colocalization of GFP and markers for parvalbumin and somatostatin was observed, demonstrating specificity of this intervention to principal neurons (Supplemental Figure 1). Furthermore, a reduction in GluN2B protein immunoreactivity was observed in cells infected with Cre-GFP, but not GFP alone, suggesting effective deletion of GluN2B (Supplemental Figure 1). Two weeks post-surgery, mice were assessed on two paradigms of depression-related behavior with good face validity and strong predictive validity for antidepressant efficacy^19–21^. Relative to GFP injected controls, Cre-injected mice lacking GluN2B in pyramidal neurons of mPFC (2BΔ^mPFC^ mice) displayed decreased immobility times on both the forced swim test (FST) and tail suspension test (TST) (Figure 1D). To control for potential confounding alterations in locomotion, mice were assessed in the open field test (OFT). Analysis revealed no alterations in locomotor activity in 2BΔ^mPFC^ mice (Figure 1E). Therefore, deletion of GluN2B from mPFC cortical pyramidal neurons is sufficient to elicit an antidepressant-like effect in adult mice in the absence of a change in locomotor behavior.

**Figure 1.**
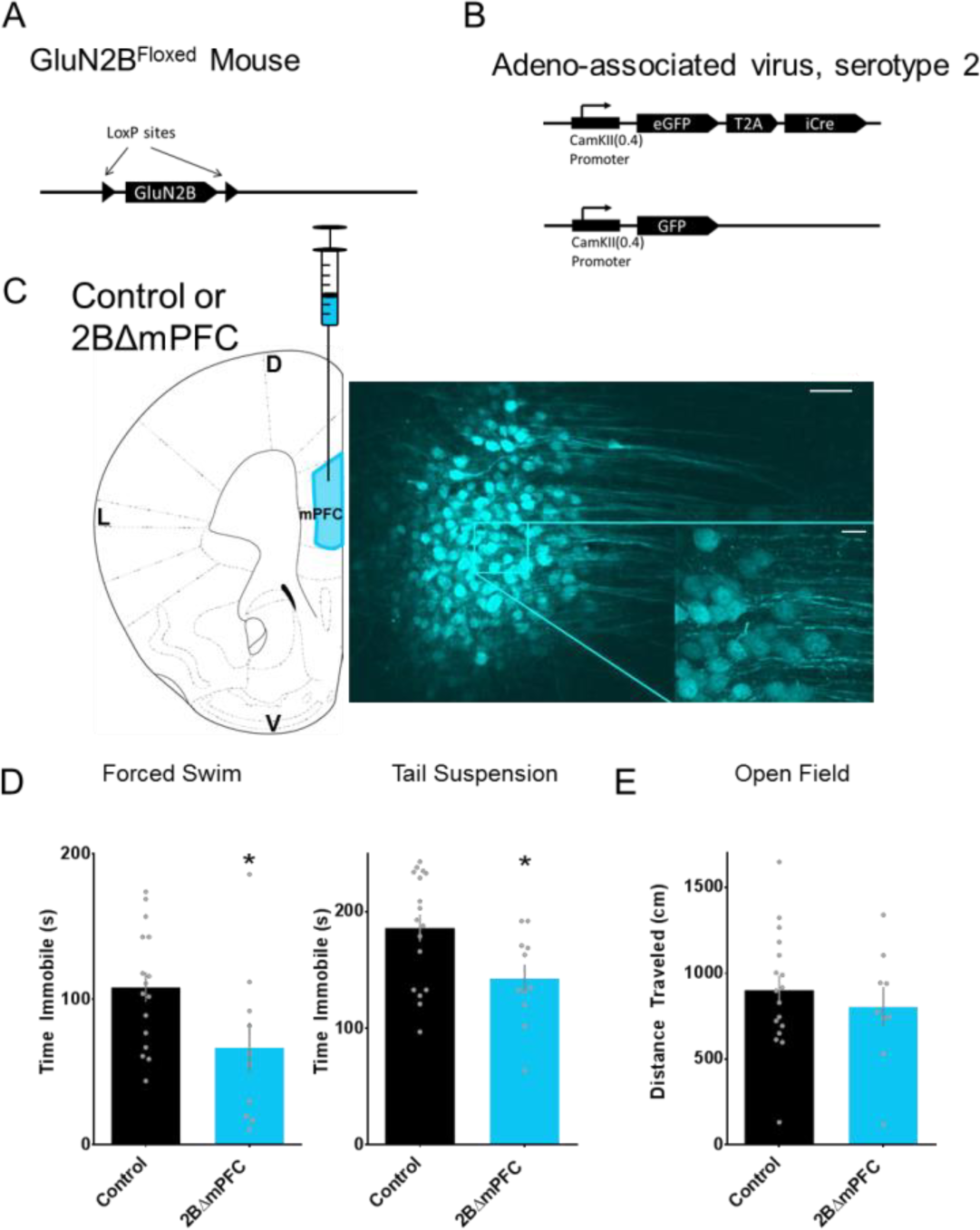
Deletion of GluN2B from mPFC pyramidal neurons elicits antidepressant-like behaviors. Deletion of GluN2B from mPFC pyramidal neurons elicits antidepressant-like behaviors (A, B) Genetic strategy for conditional deletion of GluN2B from mPFC. Mice homozygous for an allele in which exon 5 of the *Grin2b* gene is flanked by LoxP sequences were injected with AAVs driving expression of GFP (control) or GFP-Cre recombinase (2BΔ^mPFC^) under the pyramidal neuron-specific CaMKIIα promotor. (C) Homozygous GluN2B^floxed^ mice were injected with control virus or Cre into mPFC to delete GluN2B (2BΔ^mPFC^ mouse) Scale: 60µM, inset 20µM. (D) 2BΔ^mPFC^ mice displayed an antidepressant-like effect on the forced swim test and tail suspension tests (FST: Control 108.4 ± 10.18, n=16, 2BΔm^PFC^ 66.90 ± 17.12, n=10, p<0.05; TST: Control 186.0 ± 11.69, n=17, 2BΔm^PFC^: 143.0 ± 11.83, n=11, p<0.05). (E) No alteration in locomotion behavior was observed in the open field test (OFT: Control 901.2 ± 84.52, n=17, 2BΔ^mPFC^ 804.1 ± 116.2, n=9, ns)

### 2BΔ^mPFC^ neurons receive enhanced excitatory synaptic drive and display synaptic activity dependent enhanced AP firing

To identify alterations in cellular, synaptic, and circuit-level properties that might lead to the observed antidepressant effect, we first compared the action potential firing rates between control and 2BΔ^mPFC^ pyramidal neurons in layer 3 of prelimbic mPFC (Figure 2A). As cortical pyramidal neurons are quiescent at rest in acute slice, we recorded in low divalent cations (0.5mM Mg^2+^,1.0mM Ca^2+^) and injected current to depolarize neurons to −50mV holding potential. Deletion of GluN2B results in a significant increase in the rate of spontaneous action potential firing (Figure 2B) without any alteration in resting membrane potential, action potential amplitude, or half-width (data not shown).

**Figure 2.**
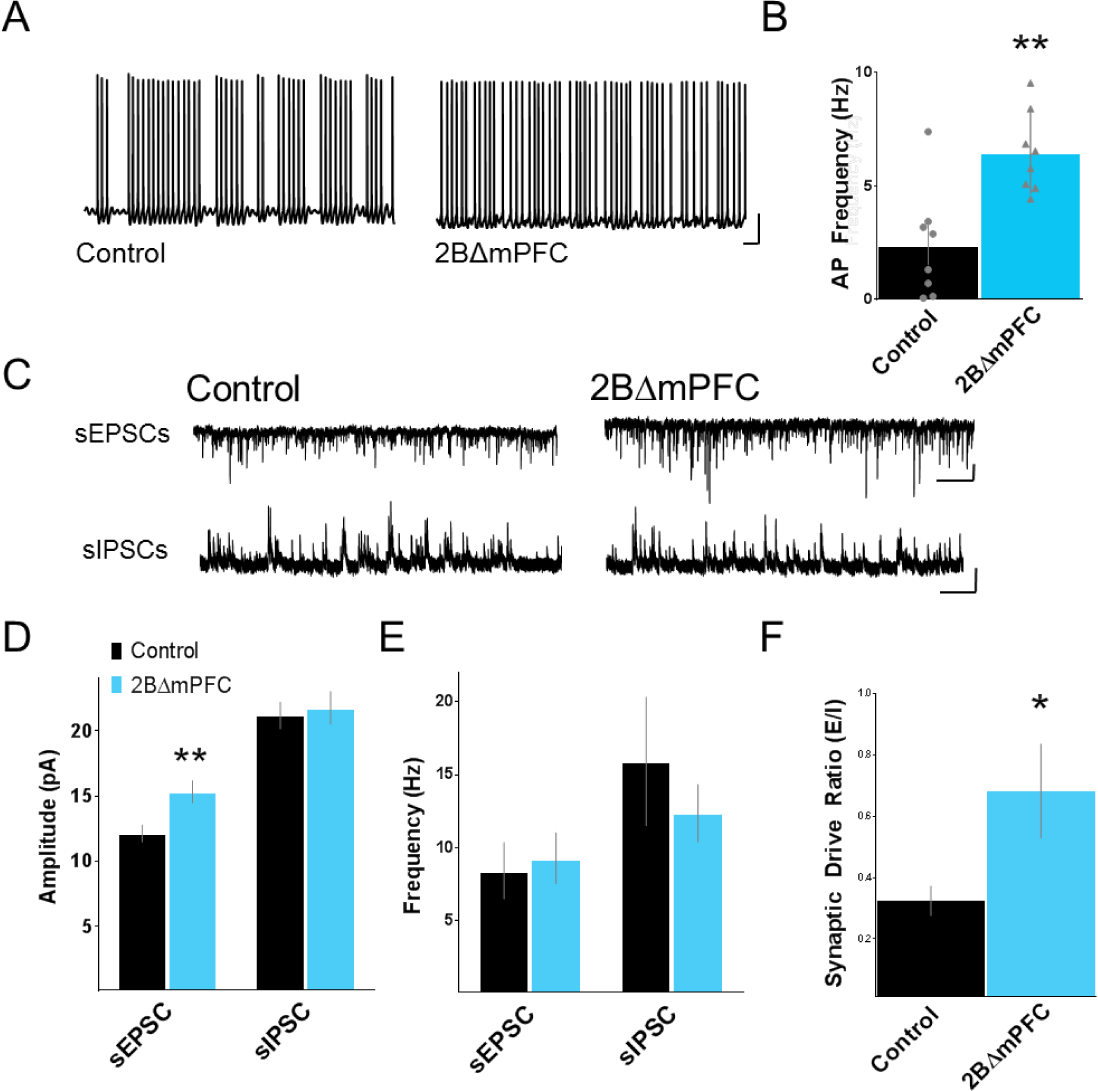
Layer 3 pyramidal 2B ΔmPFC neurons receive enhanced excitatory synaptic drive contributing to their enhanced excitability. (A) Whole-cell current-clamp recordings of control and 2BΔ^mPFC^ neurons were used to assess the frequency of spontaneous action potentials (scale: 20mV/500ms). (B) 2BΔ^mPFC^ neurons fired action potentials more readily, compared to control neurons. (Control: 2.388 ± 0.8606, n=8, 2BΔ^mPFC^ 6.440 ± 0.6316, n=8, P<0.01) (C) Voltage-clamp recordings of excitatory and inhibitory synaptic inputs onto control and 2BΔmPFC neurons (scale: 20pA/1s). (D) Analysis of EPSC amplitudes revealed that 2BΔ^mPFC^ neurons receive enhanced excitatory input (Control: 12.119530 ± 0.652682, n=15, 2BΔm^PFC^ 15.3228± 0.862796, n=15, P<0.01). No change in IPSC amplitude was observed (Control: 21.18 ± 1.03pA, n=6, 2BΔ^mPFC^ 21.77 ± 1.24, n=9, ns) (E) Analysis of EPSC and IPSC frequency revealed no significant difference between 2BΔmPFC and control cells (EPSC Control: 8.42 ± 1.93, n=15, 2BΔm^PFC^: 9.27 ± 1.75, n=15, ns; IPSC Control: 15.91 ± 4.4, n=6, 2BΔ^mPFC^: 12.36 ± 1.96, n=9, ns) (F) Synaptic drive ratio (EPSC Freq x Amp / IPSC Freq x Amp) revealed increased excitatory synaptic drive in 2BΔ^mPFC^ neurons (Control: 0.324 ± 0.048, n=7, 2BΔ^mPFC^ 0.682± 0.153, n=8, P<0.05)

Mechanisms that could underlie this increased spontaneous firing include enhanced intrinsic excitability, increased excitatory synaptic drive, or decreased inhibitory synaptic drive. We first sought to test if changes in inhibitory or excitatory synaptic drive might underlie the increased spontaneous firing rates observed in 2BΔ^mPFC^ neurons. To do so, we recorded both excitatory and inhibitory synaptic currents onto individual mPFC cells. This was accomplished by voltage-clamping cells at the GABA reversal potential (E_rev_ GABA = −68.5mV) to isolate glutamatergic EPSCs, and at the glutamate reversal potential (E_rev_ Glut = 2.5mV) in order to isolate GABAergic IPSCs (Figure 2C). Synaptic recordings revealed a significant increase in the amplitude of EPSCs in 2BΔ^mPFC^ neurons, while no change in IPSC amplitudes was observed (Figure 2D). Alternatively, we observed no significant change in the frequency of either excitatory or inhibitory PSCs (Figure 2E). Integrating excitatory and inhibitory amplitudes and frequencies to calculate an overall synaptic drive ratio^22,23^ revealed a 110.6% increase in the relative excitation experienced by 2BΔ^mPFC^ neurons (Figure 2F). These data demonstrate that GluN2B deletion enhances excitatory synaptic drive onto to layer 3 pyramidal neurons and augments their firing probability and information transfer downstream.

### Investigating intrinsic excitability of 2BΔ^mPFC^ neurons

To test for alterations in intrinsic excitability of 2BΔ^mPFC^ neurons, we injected hyperpolarizing or depolarizing current steps from - 70mV and recorded the resultant change in voltage. We observed no significant alteration in the current-voltage relationship (Supplemental Figure 2A) nor in the input-output relationship between control and 2BΔ^mPFC^ cells (Supplemental Figure 2B). Qualitative assessment of the hyperpolarizing current steps suggested that the HCN-mediated sag potential was enhanced in 2BΔ^mPFC^ neurons, although the recording conditions were not suited for quantitative analysis. Therefore, we sought to directly test for alterations in Ih that might underlie the differential excitability between conditions. HCN channels are voltage and cyclic nucleotide-gated ion channels. Hyperpolarization induces HCN channel opening resulting in a depolarizing cation conductance (Ih). Depending on cellular localization and composition of HCN channels, they can either *decrease* neuronal excitability by shunting incoming synaptic events or decreasing membrane resistance^24,25^, or they can *increase* excitability by enhancing recovery from spike after-hyperpolarization, enhancing post IPSP rebound, or by setting a depolarized resting membrane potential, according to observations made in hippocampus^26,27^, thalamus^28,29^, dorsal root ganglion^30^ and modeled in cortex^31^.

In fast synaptic isolation and with TTX and Ba^2+^ to block voltage gated sodium channels and inwardly rectifying potassium channels, respectively, we applied hyperpolarizing voltage steps from −50mV to −130mV to these neurons (Supplemental Figure 2C, top) and analyzed the resultant current responses (Supplemental Figure 2C, middle). This revealed an increase in Ih current density in 2BΔ^mPFC^ pyramidal neurons compared to control neurons, showing that removal of GluN2B enhances the functional expression of HCN channels in layer 3 cortical pyramidal neurons (Supplemental Figure 2E, F *note magenta lines highlighting the difference in I_h_ between control and 2B*Δ^*mPFC*^ *cells).* Note that blockade of HCN channels with ZD 7288 results in a loss of the delayed inward current during hyperpolarizing voltage steps, confirming that the currents assessed are I_h_ (Supplemental Figure 2C, bottom).

I_h_ is altered by differential expression of HCN isoforms (with data from expression systems showing that HCN1 is activated at relatively depolarized potentials, HCN4 at more hyperpolarized potentials, and HCN2 and 3 being intermediate^32^), and by open probability, dictated by the relative abundance of intracellular cyclic nucleotides. To test the voltage dependence of I_h_, we assessed the activation curve by analyzing the amplitude of tail currents after bringing each step to −130mV. We observed a hyperpolarizing shift in the voltage dependence of I_h_ activation in 2BΔ^mPFC^ neurons compared to control (Supplemental Figure 2G), demonstrated as a - 7.98mV shift in the V1/2 of 2BΔ^mPFC^ neurons (Supplemental Figure 2H). Because the increase in 2BΔ^mPFC^ cells’ I_h_ is observable only at potentials hyperpolarized to that at which we measured action potential firing it is unlikely that alterations in this property contributes to the enhanced spontaneous firing phenotype observed in 2BΔ^mPFC^ cells seen in Figure 2A, B. However, due to potential incomplete space clamp when using potassium-based internal solution, a role for I_h_ in this effect cannot be ruled out. The involvement of HCN channels in MDD and animal models of depression has precedence in the literature^33–35^ and requires further study.

### Identification of mPFC input nuclei

The observation that GluN2B deletion results in enhanced excitatory synaptic drive onto layer 3 pyramidal neurons led us to ask, which cortically-projecting nuclei might be responsible for sending this enhanced excitation to mPFC? Unstructured, global enhancement of synaptic input from all upstream projections would increase noise and eliminate specificity of information flow into and out from mPFC, and would presumably be detrimental to cortical processing^36–38^. We hypothesized that modification of a specific behavior (in this case depression-related behaviors, in the absence of locomotive alterations) is instead likely due to altered connectivity with a subset of connected brain regions.

The medial dorsal thalamus (MDT) is an association nucleus and the primary thalamic input to mPFC. Unlike relay nuclei in the thalamus, MDT receives no direct sensory input from the periphery. Instead, MDT integrates inputs from ventral pallidum (VP, GABAergic), substantia nigra (SN, GABAergic, dopaminergic), ventral tegmental area (VTA, dopaminergic), amygdala (Amg, glutamatergic) and dorsal raphe nucleus (DRN, undefined) and sends forward this “emotionally rich” information to mPFC^39–45^. This circuit is essential for regulation of working memory and behavioral flexibility^46–48^ and is defective in mouse models of and patients with neuropsychiatric disorders^49–51^. The ventral hippocampus (vHipp) is a spatially and functionally distinct section of the hippocampal formation^52^. It has established roles in the expression of social behavior^53^, anxiety^54–57^, and depression-related behavior^58^. In order to verify that our mPFC coordinates do in fact receive inputs form MDT and vHipp, we injected canine adenovirus type 2 expressing Cre recombinase (CAV-Cre)^59,60^ into the mPFC of the Ai14 Cre-dependent reporter mouse line^61^ to selectively label neurons that send axonal inputs into mPFC (Supplemental Figure 3A). Histological analysis confirmed direct inputs to mPFC from several brain regions including MDT, vHipp, and contralateral mPFC (Supplemental Figure 3A-D). Based on these retrograde tracing data, and their known role in neuropsychiatric disorders, we selected MDT and vHipp for electrophysiological analysis.

To selectively control these mPFC input nuclei, we injected an AAV expressing the excitatory optogenetic actuator Chronos^62^ into MDT or vHipp (Supplemental Figure 3E, F) and 3-4 weeks post-injection prepared coronal slices containing the targeted areas for patch-clamp recordings. Whole-cell recordings from neurons in MDT or vHipp confirmed large inward currents and reliable action potential generation with photostimulation of Chronos-infected neurons (Supplemental Figure 3D, F).

### Glutamatergic MDT and vHipp principal neurons are monosynaptically connected to pyramidal neurons and drive feed-forward inhibition in mPFC

To define the nature of functional connectivity of inputs into mPFC, we injected mice with Chronos in MDT or vHipp and prepared PFC slices containing Chronos^+^ axon terminals for recordings (Supplemental Figure 4A, G). GFP-labeled Chronos axons were most predominantly visible in cortical layers 1 and 3 in MDT-injected mice, and throughout cortical layers in vHipp-injected mice. Stimulation of MDT and vHipp inputs with 1ms, 473nm light pulses reliably evoked excitatory synaptic currents in layer 3 mPFC pyramidal neurons recorded at the GABAA reversal potential (Supplemental Figure 4B, H), and inhibitory synaptic currents when held at the glutamate reversal potential (Supplemental Figure 4C, I). Analysis of the time course of the synaptic responses revealed that both the MDT and vHipp’s excitatory response occurred with low latency and small variability in onset time (Supplemental Figure 4D, J) relative to the inhibitory component. Application of TTX alone abolished both excitatory and inhibitory current components, while TTX and 4-AP together recovered excitatory, but not inhibitory currents, confirming monosynaptic excitation and polysynaptic inhibition (Supplemental Figure 4E, K). Consistent with this, only inhibitory currents were abolished by GABAA antagonist picrotoxin, while both excitatory and inhibitory currents were abolished by glutamate receptor blockade with NBQX (Supplemental Figure 4F, L). Together these data suggest that inputs from both MDT and vHipp elicit monosynaptic glutamatergic input and disynaptic, feed-forward GABAergic inhibition.

### Distinct short-term plasticity of MDT and vHipp inputs into mPFC

In order to assess the potential contributions of these two inputs to the enhancement of synaptic drive after GluN2B deletion, we must first understand their basal characteristics. To do so, we voltage clamped neurons at GABA_A_R and glutamate receptor reversal potentials and stimulated fibers with 1ms, 10Hz light pulses to assess the short term plasticity of postsynaptic currents. Analysis of the waveforms of the respective *excitatory* inputs revealed distinct kinetics, with MDT exhibiting depressing inputs, characteristic of high probability of release synapse, and vHipp exhibiting facilitating inputs, characteristic of low probability of release synapse (Figure 3A,D). Similarly, analysis of light-evoked *inhibitory* currents suggests that both inputs elicit depressing short-term plasticity, although with distinct kinetics (Figure 3B,E). These alternate forms of monosynaptic glutamatergic and feed-forward GABAergic synaptic input will have distinct consequences for information transfer from these presynaptic nuclei due to their differential ability to elicit postsynaptic action potential firing in mPFC.

**Figure 3.**
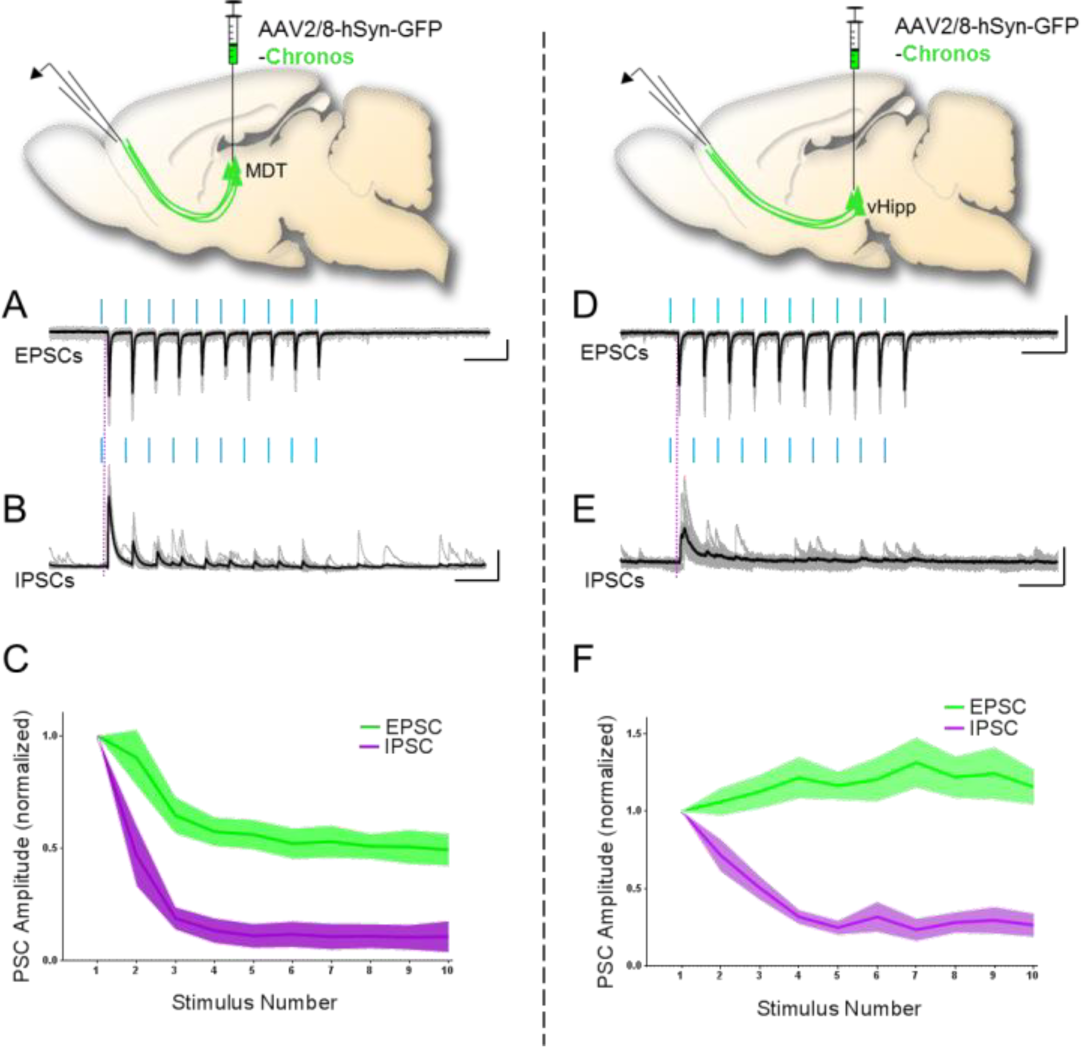
MDT and vHipp display distinct short term plasticity of glutamatergic and GABAergic inputs to layer 3 mPFC pyramidal neurons. (A, D) 10Hz trains of 1ms blue light pulses elicited robust postsynaptic EPSCs in layer 3 mPFC pyramidal neurons. MDT inputs exhibited rapidly depressing kinetics, whereas vHipp EPSCs facilitated (scale; 50pA/200ms) (B, E) 10Hz trains of 1ms blue light pulses elicited robust postsynaptic IPSCs in layer 3 mPFC pyramidal neurons. Both elicited depressing IPSCs, but with distinct kinetics (scale: B: 200pA/200ms; E: 50pA/200ms). (C, F) Summary data for short term plasticity of MDT or vHipp synaptic inputs into mPFC.

### Deletion of GluN2B from mPFC pyramidal neurons uniquely enhances MDT → mPFC inputs

Next, we investigated how these inputs are altered in the 2BΔ^mPFC^ mouse. To this end, we combined optogenetic-assisted stimulation of MDT or vHipp inputs with viral-mediated deletion of GluN2B from postsynaptic mPFC pyramidal neurons (Figure 4A, D). We observed that postsynaptic deletion of GluN2B resulted in enhancement of glutamatergic MDT inputs measured as an enhancement of the high probability of release, depressing signature (Figure 4B). In contrast, no change was observed in vHipp inputs (Figure 4E). We additionally observed that MDT-elicited feed-forward inhibition decremented more slowly in the absence of postsynaptic GluN2B (Figure 4C).

**Figure 4.**
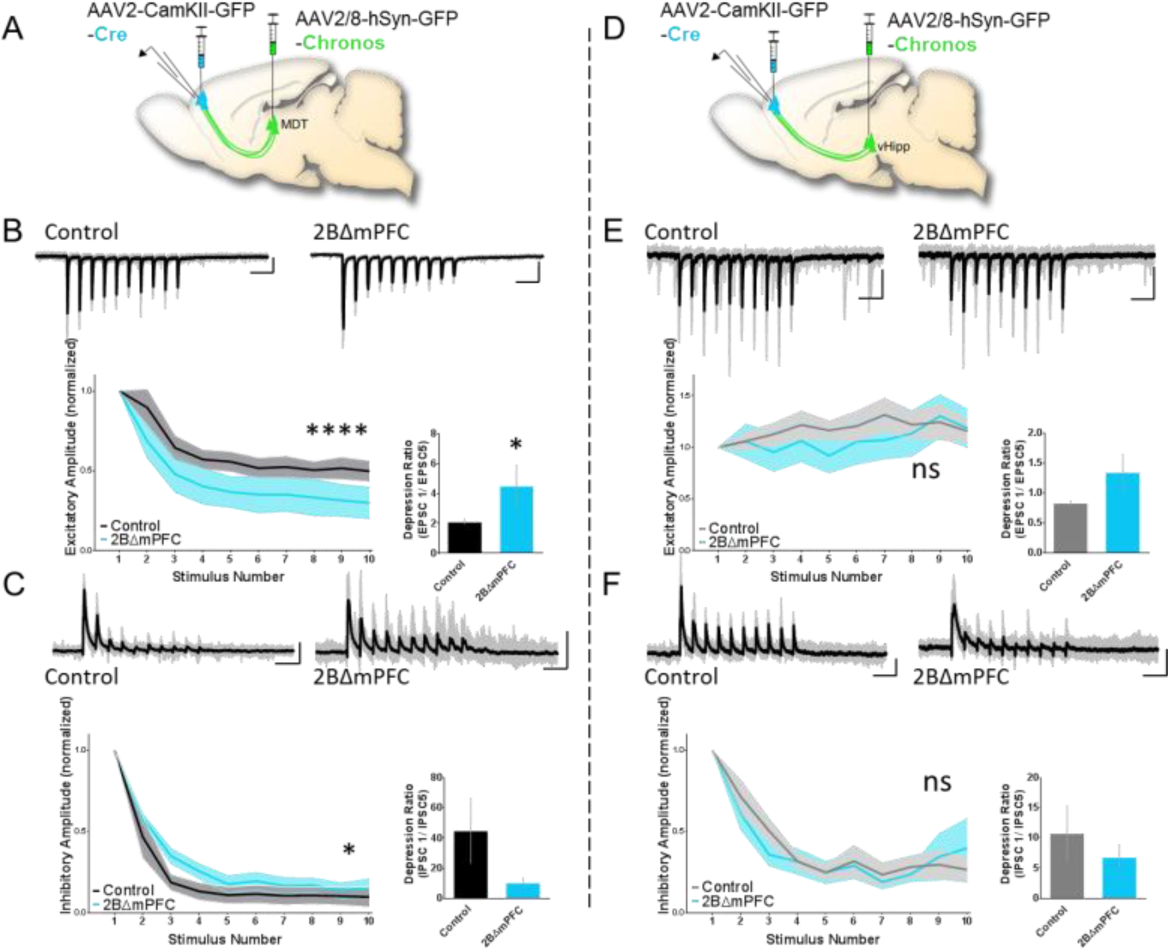
MDT→ mPFC inputs are uniquely modified by deletion of GluN2B from postsynaptic pyramidal neurons in mPFC. (A, D) Schematic of MDT experiment: Chronos was injected into MDT (left panel) or vHipp (right panel), and mPFC was injected with CaMKII-Cre to postsynaptically delete GluN2B from pyramidal neurons. Coronal sections of PFC were prepared for whole-cell recording in prelimbic mPFC and photostimulation of afferent fibers. (B, E) Postsynaptic deletion of GluN2B amplifies the high probability of release, depressing phenotype of MDT excitatory inputs into mPFC (MDT Control: n=17; scale: 40pA/200ms, 2BΔ^mPFC^: n=7, scale 100pA/200ms; two-way ANOVA F (1, 190) = 15.91, P < 0.0001). Conversely, postsynaptic deletion of GluN2B has no effect on the kinetics of vHipp excitatory inputs into mPFC (vHipp Control: n=17, scale: 20pA/200ms, 2BΔ^mPFC^: n=6, scale 20pA/200ms; two-way ANOVA F (1, 210) = 1.686, P = 0.1955. (C, F) Postsynaptic deletion of GluN2B results in a sustained feed-forward inhibitory response throughout the stimulus train in MDT inputs (MDT Control: n=17, 2BΔ^mPFC^: n=7 two-way ANOVA F(9, 110) = 16.57, P < 0.05). Conversely, postsynaptic deletion of GluN2B has no effect on the kinetics of vHipp inhibitory inputs into mPFC (vHipp Control: n=17, 2BΔ^mPFC^: n=6 two-way ANOVA F (1, 200) = 0.1980, ns, scale: E: 100pA/200ms, F: 50pA/200ms).

As the variability in expression of Chronos between animals makes direct amplitude comparisons difficult to interpret, we next recorded sequentially paired Cre^+^ (2BΔ^mPFC^) and Cre^−^ (control) cells within the same mPFC slices and made direct comparisons. Deletion of GluN2B resulted in the significant enhancement of amplitude MDT driven EPSCs, while having no effect on the amplitude of vHipp-driven EPSC amplitude (Figure 5A, B, E, F)

Finally, we tested how this enhanced MDT input onto GluN2B-null cells relates to action potential output from these neurons. In current clamp recordings from GFP control neurons, stimulus trains never resulted in action potential output in any recorded cells in response to either MDT or vHipp fiber stimulation. However, in 2BΔ^mPFC^ cells, MDT stimulation, but not vHipp stimulation was sufficient to drive spiking in 71.4% of recorded layer 3 mPFC pyramidal neurons. Moreover, paired GFP-negative cells in injected slices did not fire action potentials in response to stimulation (Figure 5C, G, D, H). Together, these data show that deletion of GluN2B from mPFC uniquely facilitates MDT inputs and allows for synaptic stimulation-induced action potential generation.

**Figure 5.**
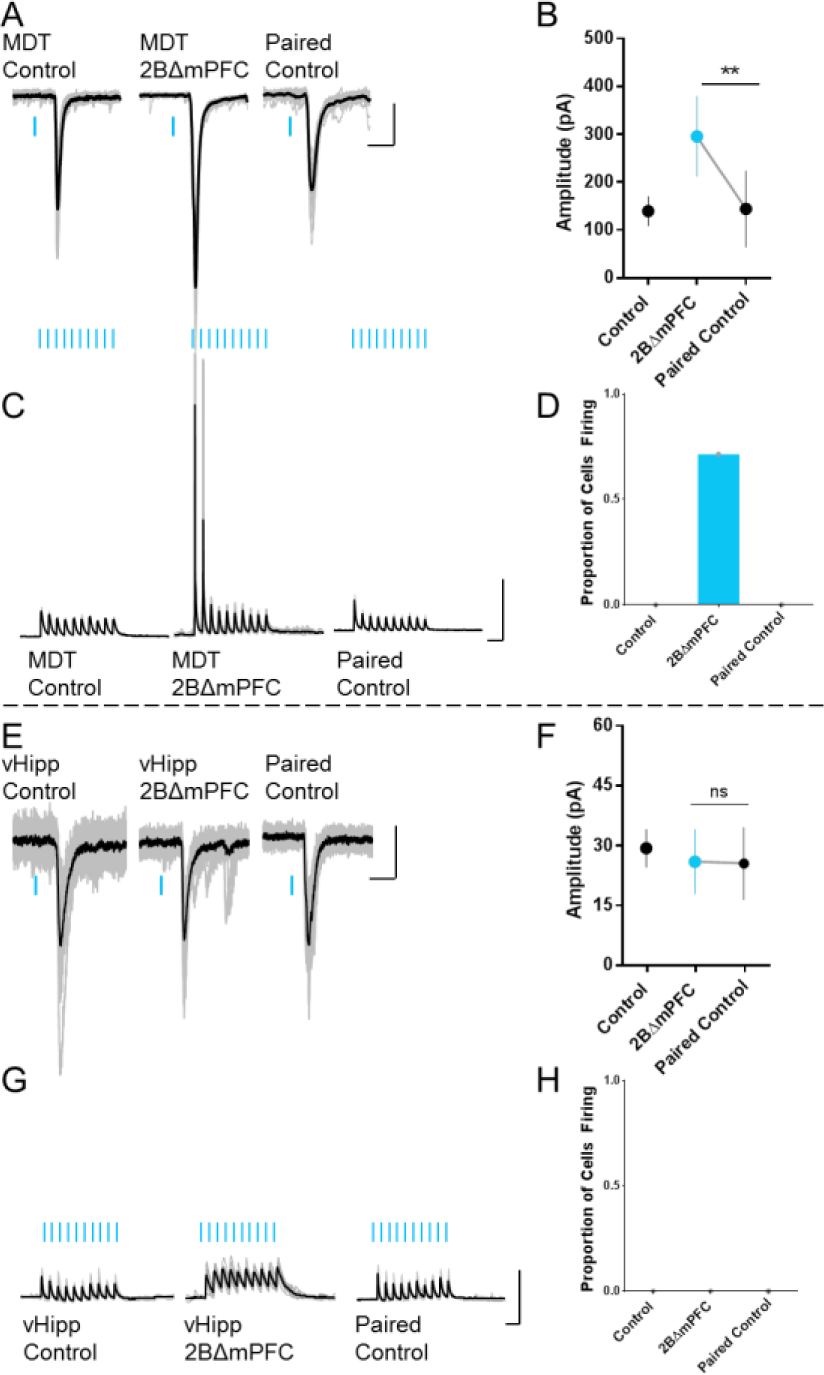
Deletion of GluN2B from mPFC pyramidal neurons increases synaptic drive and permits synaptically-driven action potential generation after MDT fiber stimulation. (A) Sample traces of EPSCs after MDT fiber stimulation in control animals, Cre+ cells from 2BΔ^mPFC^ animals, and paired Cre-cells in 2BΔ^mPFC^ animals. (B) Statistical comparison of paired control and 2BΔ^mPFC^ neurons demonstrates enhanced synaptic drive after GluN2B deletion (Paired control: 152.0 ± 35.3, n=8, 2BΔ^mPFC:^ 295.8 ± 72.57, n=8, P<0.01) (C, D) Postsynaptic deletion of GluN2B alters MDT inputs such that trains of synaptic stimulation elicit postsynaptic action potential generation in mFPC pyramidal neurons (0.0% of Control cells, 71.4% of 2BΔ^mPFC:^ cells, 0.0% of paired control cells). (E) Sample traces of EPSCs after vHipp fiber stimulation in control animals, Cre+ cells from 2BΔ^mPFC^ animals, and paired Cre-cells in 2BΔmPFC animals. (F) Statistical comparison of paired control and 2BΔ^mPFC^ neurons demonstrates no change in synaptic drive after GluN2B deletion (Paired control: 25.58 ± 6.42, n=3, 2BΔ^mPFC:^ 26.04± 5.75, n=3, ns) (G, H) Postsynaptic deletion of GluN2B does not alter firing drive of vHipp input (0.0% of Control cells, 0.0% of 2BΔ^mPFC:^ cells, 0.0% of paired control cells).

### MDT→ mPFC synaptic NMDARs are GluN2B-rich

The observation that deletion of GluN2B from mPFC pyramidal neurons results in enhancement of synaptic drive from MDT but not vHipp led us to question what possible mechanism could account for this specificity. We hypothesized that differential basal expression of GluN2B at these two synapses might provide a substrate for why MDT but not vHipp inputs are altered after GluN2B deletion. To test this hypothesis, we again recorded from layer 3 pyramidal neurons in mPFC and drove monosynaptic NMDAR-mediated synaptic currents under basal conditions and in the presence of the GluN2B-specific antagonist ifenprodil (3uM) (Figure 6A, C). Analysis revealed that GluN2B-containing NMDARs account for a significant proportion of synaptic NMDAR-mediated current at the MDT→ mPFC (34.9% of NMDAR current p<0.01) but not vHipp→ mPFC synapse (12.52% of NMDAR current, ns) (Figure 6B, D. Subsequent application of pan-NMDAR blocker APV resulted in the elimination of the remainder of the NMDAR-mediated current, confirming the specificity of our measurement (Figure 6B, D). We previously demonstrated that GluN2B-containing NMDARs suppress the functional expression of AMPA receptor-mediated current ^9,63^. Accordingly, we found that the MDT→ mPFC synapse, being rich in synaptic GluN2B, had a significantly reduced AMPA: NMDA ratio, relative to the vHipp input (Figure 6E–G). Based on these data and our previous findings, deletion of GluN2B from pyramidal neurons in mPFC would lead to AMPA receptor incorporation at synapses receiving input from MDT specifically, and might produce the observed enhancement of excitatory synaptic drive in MDT→mPFC circuits.

**Figure 6.**
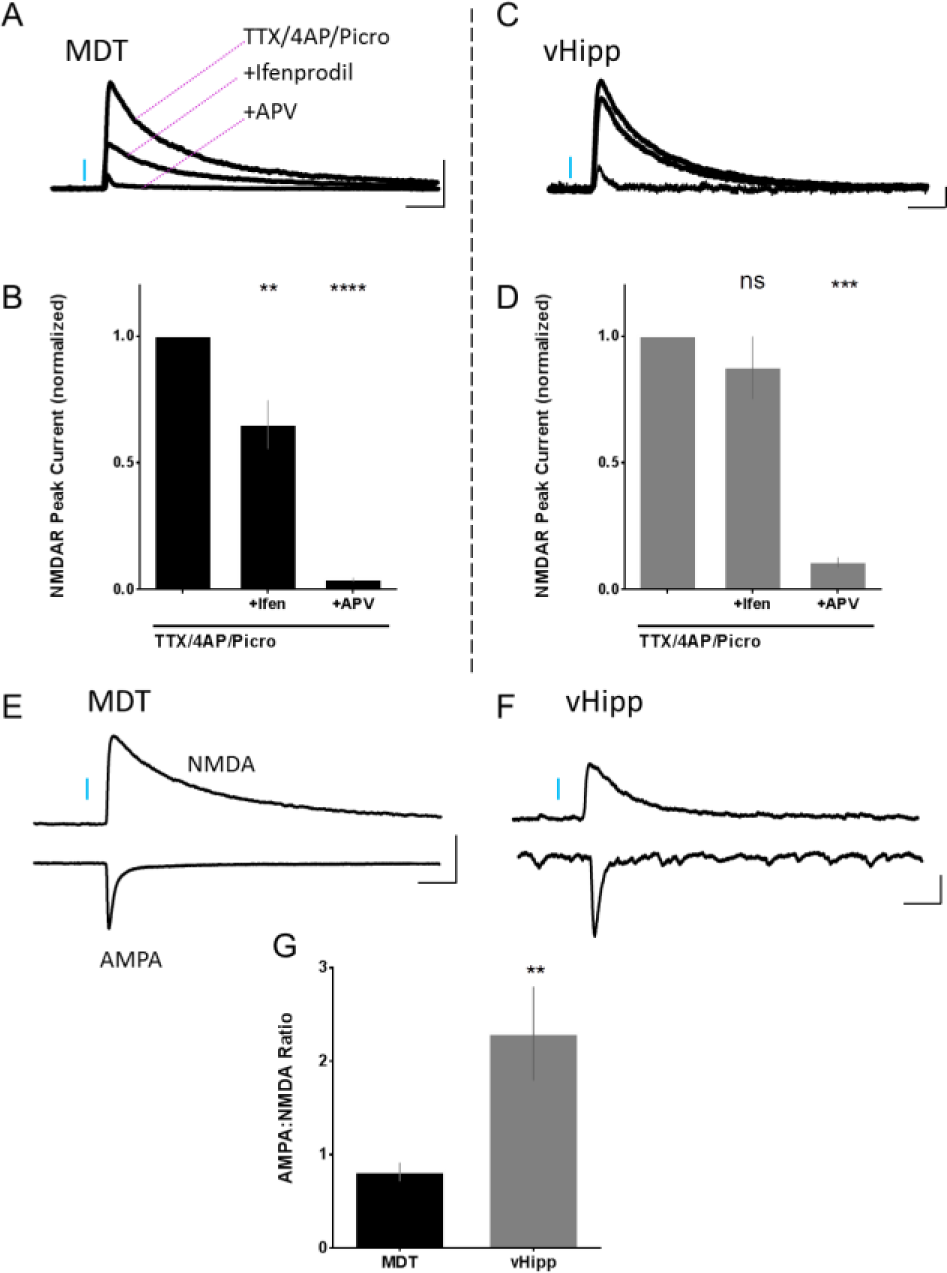
MDT→ mPFC input is rich in synaptic GluN2B. (A) Sample traces of NMDA receptor mediated current from MDT→mPFC input basally, with ifenprodil, and with APV in bath. (B) Quantification reveals that a significant proportion of the NMDA receptor mediated current (34.9%, p<0.01) is mediated my GluN2B containing NMDARs. (C) Sample traces of NMDA receptor mediated current from vHipp→mPFC input basally, with ifenprodil, and with APV in bath. (D) Quantification reveals that NMDA receptor mediated current is not significantly GluN2B mediated (12.52%, ns). (E, F) Sample traces of AMPA receptor (measured at −70mV) and NMDA receptor (measured at +40mV, measured 50ms after stimulus) mediated currents driven by MDT and vHipp inputs. (G) Quantification demonstrates that MDT inputs have a significantly lower AMPA:NMDA ratio.

### Investigation of mPFC-associated circuits using perfusion fMRI in 2BΔ^mPFC^ mice

Given the changes observed at the level of mPFC and MDT, we next sought to understand how these changes lead to altered activity in mPFC-connected nuclei *in vivo.* To determine which brain regions are altered in 2BΔ^mPFC^ mice, we injected GluN2B floxed mice with Cre virus (unilaterally) or with GFP control virus and employed a perfusion fMRI approach. Considering an *a priori* defined subset of regions of interest (ROIs) (primary endpoints) based on established mPFC connectivity and known role in neuropsychiatric disorders, perfusion analysis revealed significantly altered activity in ipsilateral mPFC (at the injection site) and MDT, but not in ipsilateral vHipp, dorsal raphe nucleus (DRN), dorsal periaqueductal grey (dPAG), or any contralateral ROI(Figure 7A). Visualization of all ROIs at a descriptive level (secondary endpoints: Figure 7B) demonstrates the brain-wide changes in perfusion, presented without statistical analysis.

**Figure 7.**
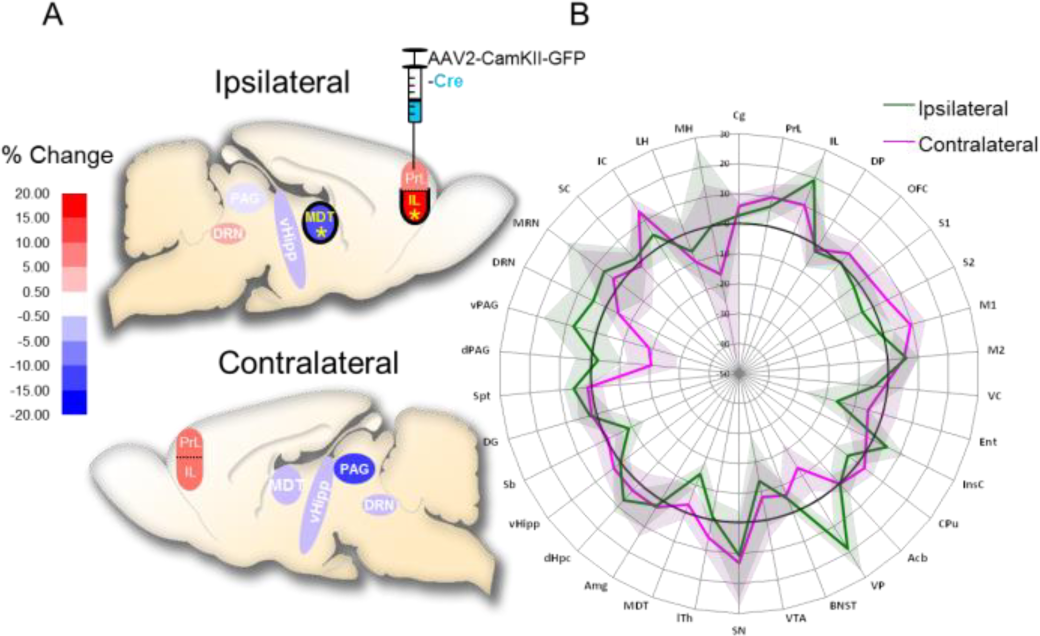
Perfusion fMRI reveals alterations in mPFC and MDT in 2BΔ^mPFC^ mice. (A) Perfusion alterations (given in % normalized perfusion) after unilateral injection of Cre-GFP virus (as compared to uni- and bilateral injection of GFP control virus) in 6 *a priori* defined ROIs. Significant alterations were seen in the mPFC and MDT of the ipsilateral hemisphere (yellow *: passing false discovery rate control at 10% across ROIs and hemispheres; based on 2-way ANOVA followed by contrasting “Cre(unilateral)” with “GFP(joint uni- & bilateral)”; GFP: n=14, Cre-GFP(unil.): n=5). (B) Radar plot. Descriptive overview of the same perfusion contrast in all ROIs available from the in-house brain atlas. Colored lines and shaded error bands indicate means and standard errors of the perfusion differences, respectively (Bold black line at zero: GFP control level).

## Discussion

Depression is a complex neuropsychiatric disorder arising from disruption of normal cellular, synaptic, and circuit functions involving the mPFC^64–67^. The NMDAR antagonist ketamine has been shown to normalize these disrupted physiological features, alter functional connectivity of mPFC, and elicit antidepressant effects in human patients and rodent models of depression^14,68,69^. Because ketamine’s antidepressant effect is mimicked and occluded by blockade of GluN2B-containing NMDA receptors in cortex, we sought to explore the behavioral, cellular, synaptic, and circuit-level consequences of post-developmental GluN2B-knockout in pyramidal neurons of mPFC. We found that this manipulation drives an antidepressant-like response in mice. This also resulted in enhanced excitatory synaptic drive onto, and increased action potential output from, these neurons. Optogenetic-assisted circuit analysis demonstrated that GluN2B deletion enhances the high probability of release phenotype of MDT→mPFC inputs, and specifically facilitates MDT synaptically-driven action potentials in pyramidal neurons of mPFC. These findings improve our understanding the role of NMDAR blockade as an antidepressant, and define a novel circuit regulating depression-like behavior in mice.

### mPFC circuits and depression-related behavior

Several brain nuclei downstream from mPFC have been shown to play a role in regulating depression-related behaviors including the dorsal raphe and lateral habenula^18^, basolateral amygdala^70^ and nucleus accumbens^71^. Conversely, little attention has been paid to the roles of afferent projections to mPFC, with the notable exception of dopaminergic inputs from VTA^72–74^. We demonstrated that MDT, but not vHipp connectivity is altered after deletion of GluN2B. Interestingly, in contrast to a recent publication that demonstrated a role for vHipp→mPFC in ketamine-mediated antidepression^58^, we found no alteration in functional connectivity between vHipp and mPFC in our model. This could represent divergent mechanisms for GluN2B-specific manipulation vs. mechanisms involving pan NMDAR antagonism. Developing our understanding of the NMDAR contributions at these individual inputs will be necessary to better understand the mechanisms at play.

### Mechanism of action

How GluN2B deletion specifically alters the post-synaptic response to MDT input is another outstanding question that must be addressed. Our previous work demonstrated that deletion of GluN2B-containing NMDARs enhances the functional expression of AMPA receptors in neuronal culture and *in vivo*^9,63^. We hypothesized that MDT axons innervate dendritic spines that contain a disproportionally large proportion of GluN2B-containing NMDARs and therefore have suppressed AMPA receptor expression. Our results demonstrating higher ifenprodil sensitivity of the MDT input support this hypothesis, as does our observation that post synaptic responses to MDT stimulation have a lower AMPA: NMDA receptor-mediated current ratio. After deletion of GluN2B, we observe a compensatory facilitation of the post synaptic response to MDT stimulation specifically. Synapse-specific differences in NMDAR subunit contribution have been observed; for example, cortically-projecting thalamic inputs preferentially synapse onto GluN2B-rich synapses on layer 1 Reelin+ interneurons^75^, as well as Schaffer Collaterals, which preferentially innervate GluN2B-containing CA1 synapses^76^.

### Diverse forms of information transfer to mPFC

Specific upstream nuclei convey distinct information to mPFC pyramidal neurons and differentially regulate behavioral state. Subcellular mapping revealed that layer 2 pyramidal neurons primarily receive input from thalamus at large spines onto distal dendrites and receive few inputs peri-somatically. Conversely, inputs from basolateral amygdala primarily synapse onto relatively smaller spines on proximal dendrites in layer 3. These differences would therefore have distinct consequences for synaptic integration in the postsynaptic cell^77^. High probability of release, depressing synapses enable postsynaptic neurons to respond to input stimulus trains more quickly, whereas low probability of release synapses require integration of inputs over expanded timescales^78^. We found that vHipp→mPFC glutamatergic inputs are weakly facilitating, whereas MDT→mPFC inputs are high probability of release and depressing. Furthermore, deletion of GluN2B from mPFC enhances the high probability of release characteristics of MDT inputs, and therefore enhances firing-coupling between MDT and mPFC. Altered MDT-mPFC connectivity has been observed in patients with treatment-resistant depression^51^. Thus, it is possible that the enhanced MDT→mPFC coupling we observed after deletion of GluN2B could underlie its antidepressant-like effect, although we have not tested a causal relationship in this study. Disrupting MDT→mPFC coupling also leads to deficits in cognitive function^47^, implying that this manipulation might be pro-cognitive, although this also remains to be tested.

### Evidence from *in vivo* imaging studies

While alterations in the MDT→mPFC input have not been implicated previously in rodent models of depression, evidence exists from human imaging studies pointing to the importance of this circuit. Deep brain stimulation (DBS) is one of the most efficacious treatments for treatment resistant depression^79^. Depressed patients administered DBS to subcallosal cingulate white matter sort into responders and non-responders. Diffusion tensor imaging studies demonstrate that responders, but not non-responders, undergo extensive white matter development in prefrontal cortex and thalamus, among other regions^80^.

Resting state fMRI (rsfMRI) is used to evaluate the regional interactions in networks across the brain. In support of our hypothesis, rsfMRI studies demonstrate that functional connectivity between dorsal anterior cingulate cortex (dACC) and MDT is impaired in MDD patients, and improvements in depressive symptoms were associated with normalization of resting state functional connectivity to healthy control levels after electroconvulsive therapy (ECT)^81^. Additionally, depressed patients with low MDT-dmPFC resting state functional connectivity were associated with better outcomes from dmPFC transcranial magnetic stimulation (TMS), and successful antidepressant response to TMS was associated with increases in MDT-dmPFC connectivity^82^.

In our mouse fMRI studies we observed that after unilateral deletion of GluN2B in mPFC pyramidal neurons, regional activity is strongly increased in ipsilateral mPFC (at the Cre virus injection site) and decreased in ipsilateral MDT. Because ASL-based perfusion fMRI cannot determine directionality of circuit effects, and because the temporal resolution does not allow for parsing of monosynaptic vs network effects, one can only speculate as to the directionality of the observed perfusion changes. Since mPFC reciprocally projects to MDT, one obvious possibility is that the increased action potential output we observe is monosynaptically affecting activity in MDT. While the sign of this proposed change does not match the results of our perfusion fMRI, directionality of fMRI signal and neuronal activity are sometimes opposing^83^. Alternatively, since changes in perfusion are affected more dramatically by alterations in synaptic release than by action potential firing^84,85^, we could be observing a polysynaptic decrease in inhibition onto MDT. One intriguing possibility is that the mPFC (glutamatergic) → nucleus accumbens (GABAergic) → ventral pallidum (VP, GABAergic) → MDT pathway is involved. The increased action potential output we observe from mPFC might therefore di-synaptically decrease the amount of inhibitory drive from VP → MDT (resulting in decreased perfusion), and therefore enhance MDT→mPFC drive. Although our a priori set of ROIs did not include VP, the descriptive radar plot we derived suggests that VP perfusion might be enhanced. However, further study will be necessary to elucidate the networks involved in the observed antidepressant effect.

### Conclusions

In order to develop translatable circuit-based therapies for depressed patients, we must obtain a better understanding of the properties of relevant circuits that are impaired and seek to understand the mechanisms by which theses impairments are normalized after antidepressant treatment. We observed that MDT→mPFC inputs are specifically enhanced after a manipulation that increases motivated behavior in mice. With this understanding, we can now delve into unveiling input-specific biological characteristics that might provide a way to target the MDT→ mPFC connection. For example, identification of unique neurotransmitter/modulator receptors or ion channels might serve as a target for pharmaceutical intervention. With the plethora of single cell and cell-type specific molecular profiling tools available, this should be a tractable method to translate circuit-based biological findings into the clinic.

## Experimental Procedures

### Animal experimentation

This study was performed in strict accordance with the recommendations in the Guide for the Care and Use of Laboratory Animals of the National Institutes of Health according to approved institutional animal care and use committee (IACUC) protocols of Tulane University, and with strict adherence to the Swiss federal ordinance on animal protection and welfare as well as according to the rules of the Association for Assessment and Accreditation of Laboratory Animal Care International (AAALAC), and with the explicit approval of the local veterinary authorities in both locations. Mice were housed on a 12/12hr light–dark cycle, with regular rodent chow and tap water available *ad libitum.* Mice were weaned at ~P30 by gender. Juveniles were group-housed to not >3 animals/cage. Male, homozygous GluN2B^floxed^ mice 2-3 months of age were used at the time of analysis for all physiology, behavior, and histology studies, with the exception of the CAV-Cre tracing studies, which were done in Ai14 mice.

#### GluN2B^Floxed^ Mice

Exon 5 of the *GR1N2B* gene is flanked by LoxP sites. Expression of Cre-recombinase results in Cre-LoxP recombination and loss of GluN2B subunit expression^86^.

#### Ai14 Mice

The Ai14 mouse (Rosa-CAG-LSL-tdTomato-WPRE: ΔNeo) has a LoxP-flanked STOP cassette preventing transcription of downstream tdTomato. Expression of Cre-recombinase results in Cre-LoxP recombination and deletion of the STOP cassette, allowing for expression of tdTomato in cells of interest.

### Viral Manipulations

For spatially and temporally defined deletion of GluN2B, homozygous GluN2B^floxed^ mice were injected with *AAV2-CaMKII(0.4)-GFP(or RFP)-T2A-iCre* or *AAV2-CaMKII(0.4)-GFP(or RFP)* (control) viruses(Vector Biolabs), bilaterally or unilaterally targeted to prelimbic mPFC (ML: +/-0.30mm, AP: 1.94mm, DV: -2.50mm, 300nl). For Chronos-assisted electrophysiology, mice were injected with *AAV2/8-hSyn-Chronos-GFP* to target medial dorsal thalamus (ML: +/−0.33mm, AP: −1.34mm, DV: −3.1mm, 200nl), or CA1 of ventral hippocampus (ML: +/−3.00mm, AP: −3.16mm, DV: −4.6mm, 300nl) (University of North Carolina Vector Core). For retrograde tracing, CAV2-CMV-Cre (Centre National de la Recherche Scientifique) ^59^ was injected into prelimbic mPFC of Ai14 mice (ML: +/−0.30mm, AP: 1.94mm, DV: −2.50mm, 100nl) (JAX Stock No.007914).

### Stereotaxic Surgery

Sterile instruments and personal protective equipment were used throughout these procedures. Mice were anaesthetized with a cocktail of fentanyl (0.05mg/kg), medetomidine (0.5mg/kg), and midazolam (5mg/kg) subcutaneously and subsequently lidocaine (2%) and bupivacaine (0.5%) locally at incision site under the scalp. Animals were maintained at 35 °C throughout the surgery and afterwards until they were ambulatory. To avoid drying of the eyes during surgery, an ocular lubricant was applied. Animals’ scalps were shaved and scrubbed with betadine. After anesthetic plane was confirmed, the mouse was placed into ear-bars on a stereotaxic frame and a single rostro-caudal incision was made to expose the skull surface. Up to four small holes were made with a dental drill, and viruses or toxins were injected into sites defined by Paxinos and Watson and Allen Brain Atlas, and then confirmed empirically. Total volume per injection site varied from 100 to 300nl, as specified in the results section. Viruses were infused through a pulled glass capillary mounted to an automated microsyringe infusion pump system (Nanoject III, Drummond Scientific Company). The capillary was kept in position for at least 10min prior to retraction. Scalps were closed using octyl/butyl cyanoacrylate topical tissue adhesive (GLUture, WPI). At the end of surgery, inflammation was suppressed with meloxicam (1mg/kg, s. c.) and anesthesia was reversed with a cocktail of buprenorphine (0.2mg/kg), atipamezol (2.5mg/kg), and flumazenil (0.5mg/kg), all administered sc. Mice were monitored daily and given additional meloxicam (1mg/kg sc) as needed. After surgery, mice were single housed and allowed 14-28 days for recovery and expression before experiments were conducted.

### Electrophysiology

#### Acute Brain slice preparation

For acute slice recordings, adult 2-3 month old mice were anesthetized with isoflurane and decapitated. The brain was rapidly removed and placed into ice-cold modified NMDG solution (composition in mM: 110 NMDG, 110 HCl, 3 KCl, 10 MgCl_2_ 6*H20, 1.1 NaH_2_PO_4_ H20, 0.5 CaCl_2_ dihydrate, 25 glucose, 3 pyruvic acid, 10 ascorbic acid, 25 NaHCO_3_, 305-310 mOsm). A vibratome (Leica VT1200S) was used to generate 350µm PFC coronal sections containing the mPFC. Slices were allowed to recover in oxygenated (95% oxygen, 5% CO2) NMDG for 30 minutes at 34°C, then transferred to a holding chamber where they were incubated in bicarbonate buffered ACSF (composition in mM: 124 NaCl, 4 KCl, 26 NaHCO_3_, 1.26 NaH_2_PO_4_, 3 MgSO_4_, 2 CaCl_2_) at room temperature for at least another 30 minutes before whole-cell recordings. ACSF solutions were bubbled with 95%O_2_/5%CO_2_ at all times to maintain consistent oxygenation and pH.

#### Whole-cell patch clamp recordings

For current clamp recordings, borosilicate glass pipettes were filled with a potassium-gluconate intracellular solution (120 KGluconate, 10 HEPES, 10 D-Sorbitol, 1 MgCl_2_ 6 H_2_0, 1 CaCl_2_, 1 EGTA, 5 Phosphocreatine disodium salt hydrate, 2 ATP disodium salt hydrate, 0.3 GTP sodium salt, pH adjusted to 7.3 and 290 mOsm). For voltage clamp recordings, with the exception of I_h_ recordings, pipettes were filled with a cesium-substituted intracellular solution (120 CsMeSo_3_, 10 HEPES, 1 MgSO_4_, 0.1 CaCl_2_, 1 EGTA, 2 KCl, 5 phosphocreatine disodium salt hydrate, 2 QX-314, 2 ATP disodium salt hydrate, 0.3 GTP sodium salt). In Figure 3A, synaptic drive ratio was calculated as:

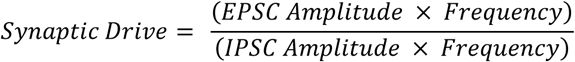

No drugs were added to bath except where otherwise noted in results section. Whole cell patch clamp recordings were acquired (MultiClamp 700B, Axon Instruments) and digitized sampled at 20kHZ (Digidata 1550, Axon Instruments), filtered at 2kHz and acquired with pClamp software (Axon Instruments). Clampfit (Axon Instruments), MiniAnalysis (Synaptosoft), and custom-written macros in IgorPRO (Wavemetrics) were used to analyze raw data and GraphPad Prism was used to perform statistical analyses, as noted in the text. Pipette resistances ranged from 2–5 MΩ. Series access resistance ranged from 3 to 14MΩ and was monitored for consistency. Cells were discarded if leak current rose above 200 pA, access resistance was greater than 15MΩ or changed by more than 15% during recording. GFP^+^ layer 3 pyramidal neurons confined to prelimbic mPFC were targeted for recording, observable with epifluorescent imaging. Spontaneous action potential activity was recorded in a modified ACSF containing 0.5mM Mg^2+^ and 1mM Ca^2+^ to closer match the composition of CSF in intact brains ^87,88^, and a small constant current was applied to set a baseline membrane potential of −50mV ^89,90^. NBQX (10µM), APV (100µM), and picrotoxin (50µM) was included in recordings in Figure 2G to achieve ionotropic synaptic isolation.

I_h_ voltage clamp experiments were recorded in NBQX (10µM), APV (100µM), and picrotoxin (50µM) to block ionotropic synaptic activity, TTX and Ba^2+^ to block voltage-gated sodium and inwardly-rectifying potassium channels, respectively. As cesium blocks HCN channels, potassium gluconate internal solution was used in these recordings. I_h_ magnitude was measured as the difference between the initial current (immediately after the capacitive transient) to the steady state current (at the end of the 2.5s voltage step). Amplitudes were normalized to individual cell capacitance to calculate I_h_ current density at a given voltage. Fractional activation of tail currents were measured relative to the largest tail current and data were fitted to a Boltzman function to determine half-activation potential:

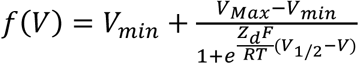

To record excitatory synaptic currents, cells were held at the measured GABA reversal potential (−68.5mV), and inhibitory currents were measured at the glutamate reversal potential (+2.5mV), corrected for liquid junction potential.

### Photostimulation of synaptic terminals

2-4 weeks after injection of 200nl AAV2/8-hSyn-Chronos-GFP into MDT or vHipp, acute brain slices were acquired as detailed above. For each experiment, the target site “hit” was visually confirmed (MDT or vHipp) and in a subset of experiments cells were patched and photostimulated to confirm functional expression of Chronos in cell bodies. PFC slices were visualized for the presence of GFP+ synaptic terminals in mPFC. Prelimbic layer 3 pyramidal neurons in the GFP+ terminal field were selected for recording. For short-term plasticity experiments 10Hz trains of 1ms pulses of blue laser light (473nm) were delivered by a galvanometer-based scanning spot illuminator (UGA-42 GEO, Rapp Optoelectronics). For paired recordings in Figure 5, neighboring Cre+ and Cre^-^ cells were pseudo-randomly selected for recording to distinguish cell-intrinsic from network effects.

### Mouse Behavior

#### Open Field Test

Open field test was conducted using Accuscan open field system. Mice were introduced into the arena at a fixed corner of an acrylic (42 × 42 × 30.5 cm) field arena and allowed to explore the arena freely. The arena was cleaned in between subjects with 70% ethanol.

#### Forced Swim Test

Forced swim test was carried out as described ^91,92^. Briefly, mice were introduced to a cylinder 15cm in diameter, 20cm height and filled to 15cm with room temperature water (23-26°C) and the mice were unable to use their tails to support themselves with the bottom of the cylinder. Two to four mice were tested simultaneously with a screen to separate the cylinders. Videos were recorded from a top-mounted camera for 6min. After testing, mice were rubbed dry and placed under an infrared lamp for ~20 min. Immobility in the last 4min of the test session was scored blind to genotype and treatment. The cylinders were cleaned in between subjects with 70% ethanol.

#### Tail Suspension Test

TST was performed as previously described ^92^. Briefly, mice were suspended from their tails using adhesive tape and video recorded for 6 min. A plastic centrifuge tube with the bottom end cut off was placed over the tail to prevent mice from climbing their tails during the test. Two to four mice were tested simultaneously with an opaque screen separating them. Videos were scored offline for immobility blind to genotype and treatment.

## Histology

Mice were transcardially perfused with 0.1 M PBS (pH 7.4) followed by 4% PFA in PBS (0.1 M, pH 7.4). Brains were postfixed in 4% PFA overnight at 4°C and then transferred to PBS and stored until ready for sectioning. Brains were sliced into 100µm coronal sections using a compresstome (Precisionary Instruments, Inc., VF300-oZ). Free-floating slices were washed 3 times for 10 minutes with PBS containing 0.5% Triton X-100 (PBST). After blocking for 2h at room temperature (RT) with PBST containing 20% normal horse serum (NHS), slices were incubated with primary anti-GFP antibody (ABCAM ab13970) in PBST containing 2% NHS at 4°C.24h later, slices were washed 3 times for 10 minutes in PBST and incubated with secondary antibody in PBST with 2% NHS for 24h at 4°C. Slices were washed again and mounted with Vectasheild Antifade Mounting Medium (Vector Laboratories H-1000). Slices were visualized on a Panoramic 250 Flash III Slide Scanner (3D Histech) with a 40X objective.

## Functional Magnetic Resonance Imaging

### Animal preparation and anesthesia

For fMRI perfusion measurements, mice were initially anesthetized with isoflurane in a carrier gas (oxygen: air 1: 5 v: v). Once animals were positioned in the MRI scanner, medetomidine anesthesia was applied by a primed continuous intravenous infusion as described previously ^93^, throughout the duration of the measurement. Animals were prone positioned with the head immobilized in a stereotaxic holder. Ophthalmic ointment was applied to the eyes to prevent corneal drying. Respiratory rate, rectal body temperature, and oxygen and CO_2_ levels in the inspired and exhaled air were continuously monitored. Body temperature was maintained at 37.3°C with a feedback-regulated electric heating blanket. After imaging, medetomidine was antagonized by a subcutaneous injection of atipamezole (Antisedan; Pfizer, Karlsruhe, Germany; 0.8 mg/kg), and animals were kept under a heating lamp until full recovery from anesthesia and then transferred back to their home cages.

### fMRI acquisition

Perfusion MRI was carried out on a Bruker Biospec 94/20 MR system (Bruker Biospin, Ettlingen, Germany). A volume resonator was used for signal excitation, and an actively decoupled quadrature surface coil was positioned over the animal’s head for signal reception. Images were acquired in 14 coronal planes between −7.10 and +4.60 mm from bregma, with an inter-plane spacing of 0.9 mm, a slice thickness of 0.6 mm and a field of view of 20 × 20 mm^2^. The first volume was a set of T2-weighted structural RARE images (TR/TEeff = 3,150/34 ms, RARE factor 8, matrix 256 × 256), followed by a T1 image series using an inversion-recovery snapshot FLASH sequence with eight inversion times (TR/TE = 4,000/1.6 ms, matrix 64 × 64) ^94^. Finally, cerebral perfusion was assessed by continuous arterial spin-labelling (CASL)^95^ with single-slice centered-RARE readout (TR/TE = 3,750 ms/5.4ms, RARE factor = 32, matrix 128 × 64, labelling pulse 3 s, post labelling delay 0.4 s, acquisition time per volume 4 min). Three successive CASL volumes were acquired and averaged for each animal.

### fMRI data analysis

Both acquisition and analysis of images were performed using an automated pipeline, thus excluding user bias. Images were processed and analyzed using in-house software written in IDL 6.4 (Interactive Data Language; Exelis, Boulder, CO, USA) and MATLAB 7.14 (The MathWorks Inc., Natick, MA, USA). Individual volumes were spatially normalized (affine plus nonlinear transform) to an in-house brain template/atlas pair, using SPM5 (Wellcome Trust Centre for Neuroimaging, London, UK). The atlas was adapted from that of Paxinos & Franklin (2001), bilaterally delineating 30+ selected anatomical brain areas as regions of interest (ROIs). T1 maps were calculated by voxel-wise fitting of a 3-parameter exponential to the T1 image series. These maps were then combined with the pertinent perfusion-weighted images to obtain quantitative perfusion maps, as described elsewhere ^96^. In order to elucidate local changes in cerebral perfusion whilst accounting for possible systemic changes affecting global brain perfusion, and to eliminate part of the inter-individual variability, individual perfusion maps were normalized slice-wise to the brain-mean value. Mean (normalized) perfusion was then determined per animal for the entire brain and within each ROI. For statistical analysis, a 2-way ANOVA was performed per ROI and hemisphere, with between-subject factors “virus type” (GFP or Cre-GFP) and “injection mode” (bilateral or unilateral). Within this model, the groups “GFP (bilateral)” (*n* = 10) and “GFP (unilateral)” (*n* = 4) were jointly contrasted with the group “Cre-GFP (unilateral)” (*n* = 5). As a primary readout, the false discovery rate (FDR) was controlled at 10 % across a subset of 6 (× 2) ROIs defined *a priori* based on prior biological knowledge. In a more exploratory way, the same contrast was reported for the remaining ROIs at a descriptive level.

## Author Contributions

O. H. M and B. J. H designed all experiments with the exception of fMRI.

O. H. M and B. J. H and wrote the manuscript, with input from A. B. and T. M.

T. M. and A. B. designed and analyzed the fMRI experiment.

O. H. M performed all other experiments and analyses, except I. B. A performed some immunohistochemical experiments.

O. H. M., A. B., T. M. and B. J. H are employees of F. Hoffmann-La Roche Ltd., Basel, Switzerland.

## Acknowledgments

We would like to thank Stephanie Schöppenthau and Sébastien Debilly for MRI data acquisition, Thomas Bielser for MRI data management and pre-processing, and Vanessa De Barros for animal colony maintenance. We also thank Eric Kremer and CNRS for providing the CAV2-Cre virus, and the Hall Lab for their invaluable comments on the manuscript.

## Supplemental Figures

**Supplemental Figure 1.**
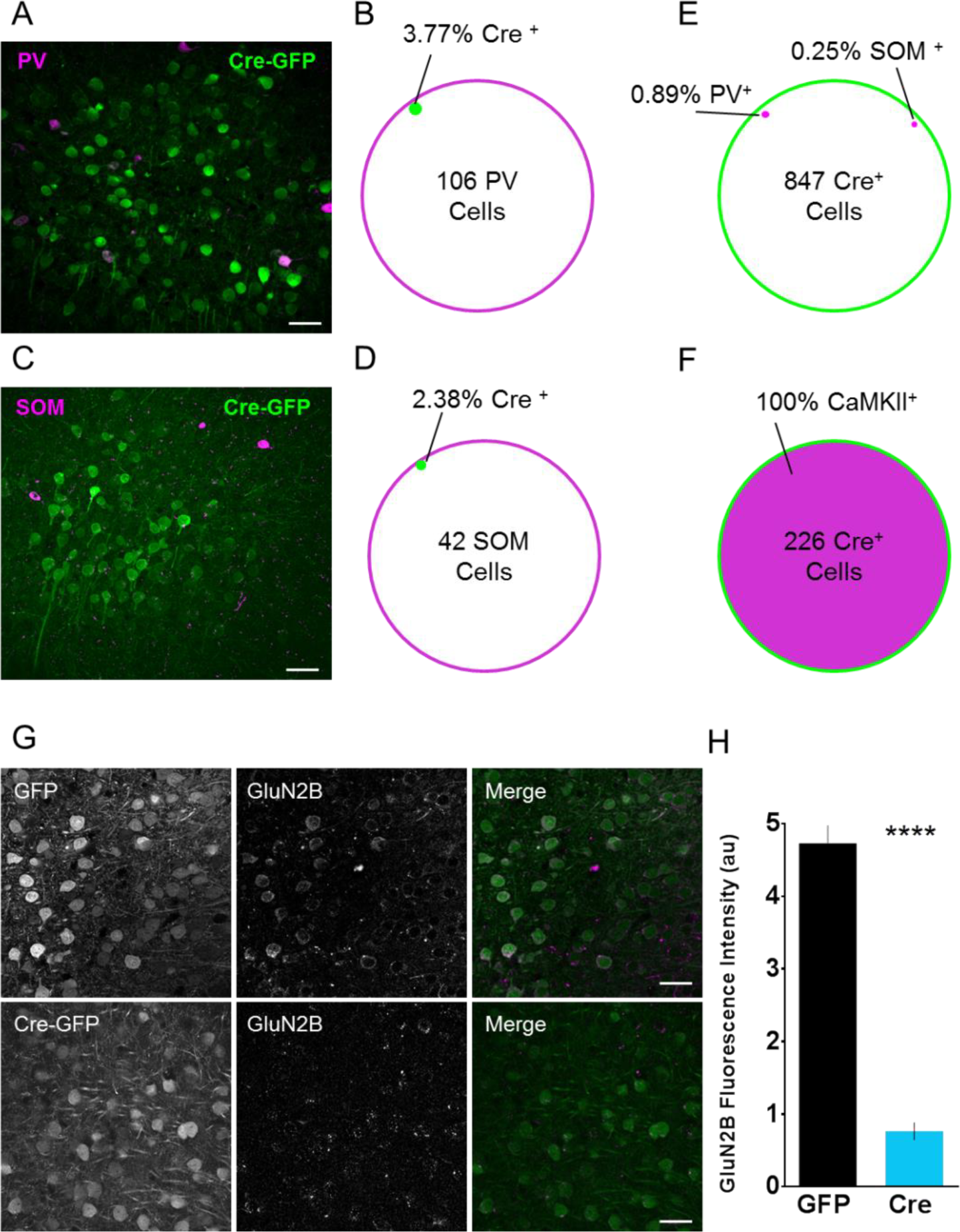
AAV-CaMKII-GFP-T2A-Cre virus is specific to pyramidal neurons and deletes GluN2B from mPFC pyramidal neurons. (A) Injection of AAV-CaMKII-GFP-T2A-Cre into mPFC results in green fluorescence in putative pyramidal neurons. Immunostaining for inhibitory interneuronal marker parvalbumin (PV) reveals minimal colocalization (Scale 60µm) (B) 3.77% of PV+ cells colocalize with virally infected cells, indicating that the virus negligibly infects PV neurons. (C) Injection of AAV-CaMKII-GFP-T2A-Cre into mPFC results in green fluorescence in putative pyramidal neurons. Immunostaining for inhibitory interneuronal marker somatostatin (SOM) reveals minimal colocalization (Scale 60µm). (D) 2.38% of SOM+ cells colocalize with virally infected cells, indicating that the virus negligibly infects SOM neurons. (E) Of the 874 putative pyramidal neurons analyzed, 0.89% colocalized with PV, and 0.25% colocalized with SOM, indicating that the virus primarily infects pyramidal neurons. (F) Of the 226 Cre+ cells assessed 100% were CaMKII+. (G, H) Injection of control AAV-CaMKII-GFP into mPFC results in green fluorescence in pyramidal neurons. Immunostaining for GluN2B protein reveals significant overlap. In contrast, injection with AAV-CaMKII-GFP-T2A-Cre significantly decreases GluN2B expression (Control: 6.496 ± 0.2456, n=197, 2BΔ^mPFC^ 2.513 ± 0.1151, n=153, P<0.0001).

**Supplemental Figure 2.**
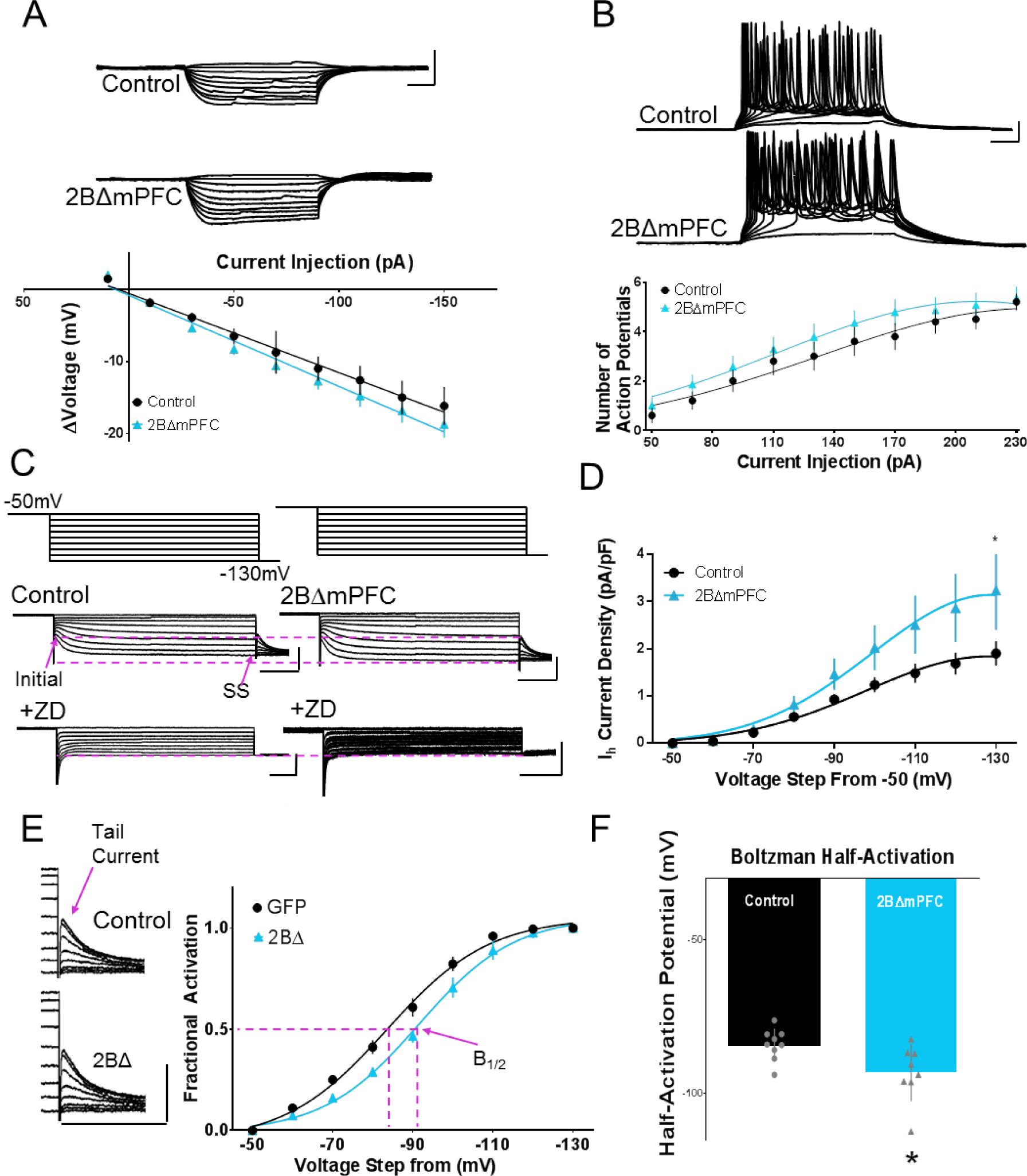
Increased spontaneous action potential output from layer 3 pyramidal 2BΔ^mPFC^ neurons is independent of alterations in intrinsic excitability. (A) The current-voltage relationship is unchanged between control and 2BΔmPFC neurons (Control: n=8, 2BΔ^mPFC^: n=13, Two-way Repeated Measures ANOVA p>0.05; scale: 20mV/40ms). (B) Action potential firing after brief depolarizing current injections is unchanged in 2BΔ^mPFC^ neurons (Control: n=10. 2BΔ^mPFC^: n= 14, Two-way Repeated Measures ANOVA P>0.05; scale: 20mV/40ms). (C) (Top) Waveform of voltage-clamp protocol: from an initial holding potential of -50mV, cells were stepped in - 10mV increments from -50 to -130mV for 2.5 seconds then each sweep was stepped to a final holding potential of -130mV to elicit a tail current. (Middle) Current responses to hyperpolarizing voltage steps recorded from control and 2BΔmPFC neurons. I_h_ was measured as the difference between initial and steady state (SS) current amplitudes (see magenta arrows). (Bottom) The HCN channel blocker ZD 7288 blocked the I_h_ current in both genotypes (Scale: 200pA/500ms (top), 100pA/500ms (bottom)). (D) Plotted values of the I_h_ current density at each hyperpolarizing voltage step revealed increased I_h_ 2BΔ^mPFC^ neurons. Data points were fit with a Gaussian curve. (Control n= 11, 2BΔ^mPFC^ n=8 Two-way Repeated Measures ANOVA Interaction F (8, 136) = 3.104, p < 0.01). (E)(Left)Sample traces showing tail currents from control and 2BΔ^mPFC^ neurons. (Right) Analysis of tail currents from voltage clamp recordings reveals a hyperpolarizing shift in the voltage dependence of the I_h_ (Scale: 200pA/500ms). (F) Fitting the data to a Boltzman function reveals 2BΔmPFC neurons are ½ activated at 7.9757mV hyperpolarized to control neurons (Control: -84.03 ± 1.706, n=9, 2BΔ^mPFC:^ -93.15 ± 3.244, n=8, P<0.05)

**Supplemental Figure 3.**
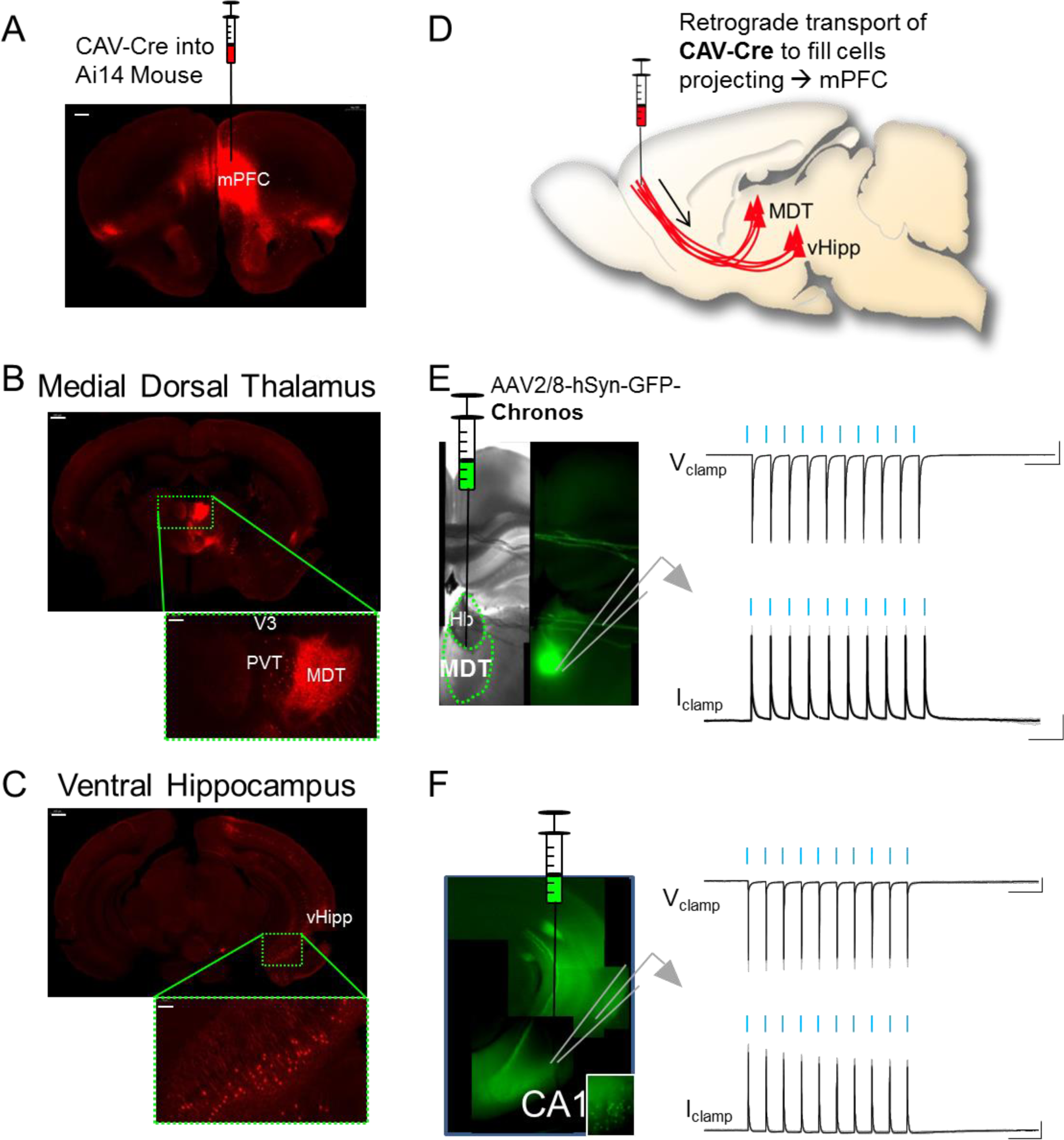
Retrograde tracing reveals synaptic inputs to mPFC from Medial Dorsal Thalamus and Ventral Hippocampus. (A) Canine adenovirus expressing Cre recombinase (CAV-Cre) unilaterally injected into ventral mPFC of a Cre-dependent reporter mouse line *(scale=500µm)*. (B) Retrograde tracing reveals a population of projection neurons in MDT that project to mPFC *(scale=500µm)*. Inset: Individual neurons visible amongst dense neurite web in MDT *(scale=100µm)*. (C) Retrograde tracing reveals a population of projection neurons in the stratum pyramidale of CA1 of the Ventral Hippocampus that project to the mPFC *(scale=500µm)*. Inset: Individual pyramidal neurons are visible in vHipp *(scale=100µm)*. (D) CAV is taken up by synaptic terminals and transported retrogradely to the cell body where Cre-LoxP recombination occurs and turns on fluorophore expression. Cell bodies of neurons projecting to mPFC are labeled with fluorophore tdTomato. (E) Injection of Chronos-GFP into MDT is visible with epifluorescent microscopy. Chronos+ cells were patch clamped for electrophysiological analysis. MDT neurons display robust photocurrents and reliable action potentials in response to 1ms, 10Hz trains of blue light (scale: 500pA/200ms; 20mV/200ms). (F) Injection of Chronos-GFP into vHipp is visible with epifluorescent microscopy. Chronos+ cells were patch clamped for electrophysiological analysis. vHipp neurons display robust photocurrents and reliable action potentials in response to 1m, 10Hz trains of blue light (scale: 200pA/200ms; 10mV/200ms).

**Supplemental Figure 4.**
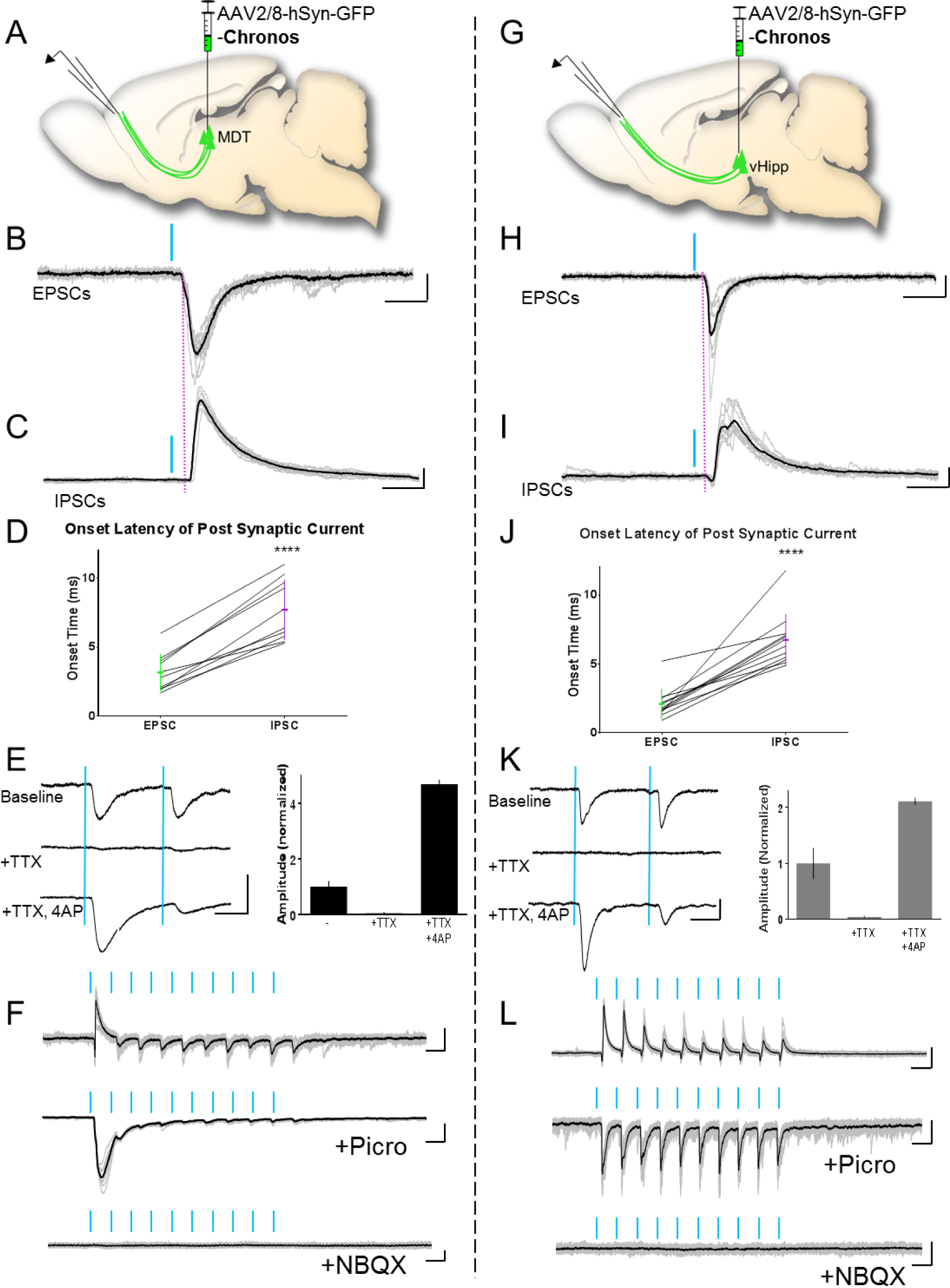
MDT and vHipp send monosynaptic glutamatergic and disynaptic GABAergic currents to layer 3 mPFC pyramidal neurons. (A, G) Schematic of MDT experiment: 3-5 weeks after Chronos injection into MDT or vHipp, coronal sections of PFC were prepared for whole-cell recording in prelimbic mPFC and photostimulation of afferent fibers. (B, H) 1ms blue light pulses elicited robust post-synaptic EPSCs in mPFC pyramidal neurons (scale: 20pA/20ms). (C, I) 1ms blue light pulses elicited robust post-synaptic IPSCs in mPFC pyramidal neurons (scale: 50pA/20ms). (D, J) Onset latency of EPSC is on the timescale of a monosynaptic input, whereas IPSC latency is on the timescale of a disynaptic input (D: n=10, p<0.001, J: n=12, p<0.001). (E, K) Left: The MDT→ mPFC and vHipp → mPFC excitatory inputs are monosynaptic, as blockade of network activity (with TTX) and stimulation of photocurrent-induced calcium flux (with 4AP) is sufficient to elicit an EPSC. Right: Sample of a single cell comparing sweeps 10 sweeps at baseline, 10 sweeps with TTX, and 10 sweeps with TTX/4AP (scale: 20pA/40ms). (F, L) Top: Voltage clamp recordings in mPFC pyramidal neurons between glutamate and GABA reversal potentials (-40mV) and light-stimulation of MDT or vHipp fibers allow for the observation of fast excitatory, and delayed inhibitory currents in the same recording. Middle: Wash in of picrotoxin (50uM) eliminates the inhibitory synaptic currents, suggesting the disynaptic component is GABAergic. Bottom: Subsequent wash in of NBQX (10uM) eliminates the excitatory component of both inputs, suggesting the monosynaptic input is glutamatergic (scale: F: 20pA/200ms (top); 50pA/200ms (middle); 10pA/200ms (bottom); L: 100pA/100ms (top); 20pA/100ms (bottom; middle)).

